# Deep neural networks and visuo-semantic models explain complementary components of human ventral-stream representational dynamics

**DOI:** 10.1101/2021.10.25.465583

**Authors:** Kamila M Jozwik, Tim C Kietzmann, Radoslaw M Cichy, Nikolaus Kriegeskorte, Marieke Mur

## Abstract

Deep neural networks (DNNs) are promising models of the cortical computations supporting human object recognition. However, despite their ability to explain a significant portion of variance in neural data, the agreement between models and brain representational dynamics is far from perfect. We address this issue by asking which representational features are currently unaccounted for in neural timeseries data, estimated for multiple areas of the ventral stream via source-reconstructed magnetoencephalography (MEG) data acquired in human participants (9 females, 6 males) during object viewing. We focus on the ability of visuo-semantic models, consisting of human-generated labels of object features and categories, to explain variance beyond the explanatory power of DNNs alone. We report a gradual reversal in the relative importance of DNN versus visuo-semantic features as ventral-stream object representations unfold over space and time. While lower-level visual areas are better explained by DNN features, especially during the early phase of the response (*<* 128 ms after stimulus onset), higher-level cortical dynamics are best accounted for by visuo-semantic features during a later time window (starting 146 ms after stimulus onset). Among the visuo-semantic features, object parts and basic categories drive the advantage over DNNs. These results show that a significant component of the variance unexplained by DNNs in higher-level cortical dynamics is structured, and can be explained by readily nameable aspects of the objects. We conclude that current DNNs fail to fully capture dynamic representations in higher-level human visual cortex and suggest a path toward more accurate models of ventral stream computations.

**SIGNIFICANCE STATEMENT:** When we view objects such as faces and cars in our visual environment, their neural representations dynamically unfold over time at a millisecond scale. These dynamics reflect the cortical computations that support fast and robust object recognition. Deep neural networks (DNNs) have emerged as a promising framework for modeling these computations but cannot yet fully account for the neural dynamics. Using magnetoencephalography data acquired in human observers during object viewing, we show that readily nameable aspects of objects, such as “eye”, “wheel”, and “face”, can account for variance in the neural dynamics over and above DNNs. These findings suggest that DNNs and humans may in part rely on different object features for visual recognition and provide guidelines for model improvement.

## Introduction

When we view objects in our visual environment, the neural representation of these objects dynamically unfolds over time across the cortical hierarchy of the ventral visual stream. In brain recordings from both humans and nonhuman primates, this dynamic representational unfolding can be quantified from neural population activity, showing a staggered emergence of ecologically relevant object information such as facial features, followed by object categories, and then the individuation of these inputs into specific exemplars (1–10). These neural reverberations are thought to reflect the cortical computations that support object recognition.

Deep neural networks (DNNs) have recently emerged as a promising computational framework for modeling these cortical computations (11). DNNs explain significant amounts of variance in neural data obtained from visual cortex in both humans and nonhuman primates (12–22). The progression of object representations from shallower to deeper layers of DNNs roughly matches the progression from lower-to higher-level object representations measured in visual cortex as neural responses unfold over time and space (13–17, 21). At the same time, DNNs also leave substantial amounts of variance in brain responses unexplained (18, 20, 23). More recent DNNs that incorporate dynamics through recurrent processing provide additional explanatory power, possibly by better approximating the dynamic computations that the brain relies on for perceptual inference (10, 24–31). However, even the most state-of-the-art DNNs still leave a significant amount of variance in brain representations unexplained (10, 20), and differences among task-trained feedforward architectures are small (32, 33), even after training and fitting (22). This raises the question of what representational features are left unaccounted for in the dynamic neural data.

To address this question, we enriched our modeling strategy with visuo-semantic object information. By “visuo-semantic”, we mean nameable properties of visual objects. Our visuo-semantic models consist of object labels generated by human observers, describing lower-level object features such as “green”, higher-level object features such as “eye”, and categories such as “face”. These object properties explain significant amounts of response variance in higher-level primate visual cortex (17, 34–45). Moreover, visuo-semantic models outperform DNNs (AlexNet (46) and VGG (47) architectures) at predicting perceived object similarity in humans (48). In addition, a recent fMRI study showed that combining DNNs with a semantic feature model is beneficial for explaining visual object representations at advanced processing stages of the ventral visual stream, especially perirhinal cortex (49). Given these findings, we hypothesized that visuo-semantic models capture representational features in ventral-stream neural dynamics that DNNs fail to account for.

We tested this hypothesis on temporally resolved magnetoencephalography (MEG) data, which can capture representational dynamics at a millisecond timescale. Human brain data acquired at this rapid sampling rate provide rich information about temporal dynamics, and by extension, about the underlying neural computations. For example, in a MEG study examining a visual delayed-match-to-sample task, time series analyses revealed distinct representational states over cue, delay, and response periods within individual experimental trials (9). In another MEG study that used source reconstruction to localize time series to distinct areas of the ventral stream, time series analyses revealed temporal inter-dependencies between areas suggestive of recurrent information processing (10).

In this work, we used representational similarity analysis to test both DNNs and visuo-semantic models for their ability to explain the representational dynamics observed across multiple ventral stream areas in the human brain. As DNNs, we used feedforward CORnet-Z and locally recurrent CORnet-R, which are inspired by the visual hierarchy of monkey visual cortex (27). As visuo-semantic models, we used existing human-generated object labels of colors, textures, shapes, object parts, subordinate categories, basic categories, and superordinate categories (45). We analyzed previously published source-reconstructed MEG data acquired in healthy human participants while they were viewing object images from a range of categories (6, 10). We investigated three distinct stages of processing in the ventral cortical hierarchy: lower-level visual areas V1-3, intermediate visual areas V4t/LO, and higher-level visual areas IT/PHC. At each stage of processing, we tested both model classes for their ability to explain variance in the temporally evolving representations. This strategy allowed us to test what visuo-semantic object information is unaccounted for by DNNs where and when during ventral-stream processing.

Our modeling strategy revealed that DNNs and visuo-semantic models explain complementary components of ventral-stream representational dynamics. DNNs outperformed visuo-semantic models in lower-level visual cortex, especially during the early phases of the response (*<* 128 ms after stimulus onset), but failed to capture a significant component of representational variance in higher-level visual cortex, for a prolonged time window starting at 146 ms after stimulus onset. This variance was accounted for by visuo-semantic labels of object parts and basic categories. For intermediate visual cortex, the explanatory power of the two model classes did not significantly differ, but the unique contribution of the DNNs peaked before that of the visuo-semantic models (around 100 versus 150 ms, respectively). The visuo-semantic peak occurred just after a period of feedback information flow from higher-level to intermediate visual cortex. Our findings point toward priorities for future modeling efforts, which should move to better capturing the dynamic computations that give rise to the emergence of visuo-semantic features represented in higher-level human visual cortex.

## Results

### DNNs better explain lower-level visual representations, visuo-semantic models better explain higher-level visual representations

We first evaluated the overall ability of the DNN and visuo-semantic models to explain the time course of information processing along the human ventral visual stream. We hypothesized that visuo-semantic models capture representational features in neural data that DNNs may fail to account for. Figure 1 shows an overview of our approach. We computed representational dissimilarity matrix (RDM) movies from the source-reconstructed MEG data to characterize how the ventral-stream object representations evolved over time in each participant. We computed a RDM movie for each participant and region of interest (ROI) and explained variance in the movies using a DNN model and a visuo-semantic model. The DNN model consisted of internal object representations in layers of CORnet-Z, a purely feedforward model, and CORnet-R, a locally recurrent variant (27), to account for both feedforward and locally recurrent computations. The visuo-semantic model consisted of human-generated labels of object features (e.g., “brown”, “furry”, “round”, “ear”; 119 labels) and categories (e.g., “great dane”, “dog”, “organism”; 110 labels) for the object images presented during the MEG experiment (45). We computed model predictions by linearly combining either all DNN layers or all visuo-semantic labels to best explain variance in the RDM movies across time. We evaluated the model predictions on data for images left out during fitting. For each model, we tested if and when the variance explained in the RDM movies exceeded the prestimulus baseline using a one-sided Wilcoxon signed-rank test. We also tested if and when the amounts of explained variance differed between the two models using a two-sided Wilcoxon signed-rank test. We controlled the expected false discovery rate at 0.05 across time points.

**Fig. 1.**
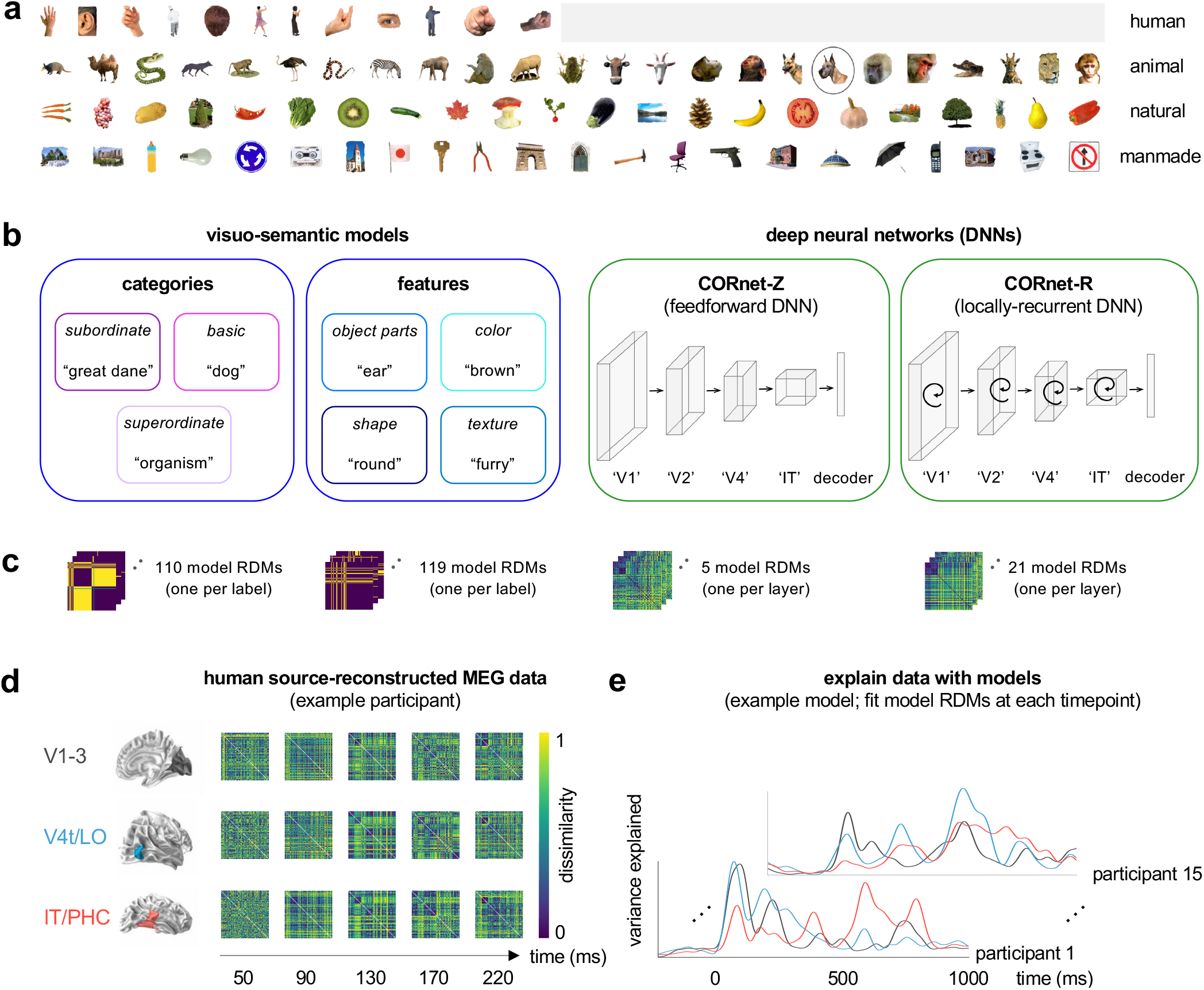
Schematic overview of approach: stimulus set, models, data, and model fitting. a) Stimulus set. Stimuli are 92 colored images of real-world objects spanning a range of categories, including humans, non-human animals, natural objects, and manmade objects (human face images are not shown and are replaced by gray squares according to bioRxiv’s policy on posting images of individuals). **b)** Visuo-semantic models and deep neural networks (DNNs). Visuo-semantic models consist of human-generated labels of object features and categories for the 92 images. Example labels are shown for the dog face encircled in panel (a). DNNs are feedforward and locally recurrent CORnet architectures trained with category supervision on ILSVRC. These architectures are inspired by the processing stages of the primate ventral visual stream: from V1 to inferior temporal cortex (IT). **c)** Object representations for each model. We characterized object representations by computing representational dissimilarity matrices (RDMs). We computed one RDM per model dimension, i.e. one for each visuo-semantic label or DNN layer. For each visuo-semantic model dimension, RDMs were computed by extracting the value for each image on that dimension and computing pairwise dissimilarities (squared difference) between the values. For each CORnet-Z and CORnet-R layer, RDMs were computed by extracting an activity pattern across model units for each image and computing pairwise dissimilarities (1 minus Spearman’s r) between the activity patterns. **d)** Human source-reconstructed MEG data for an example participant. MEG data were acquired in 15 healthy adult human participants while they were viewing the 92 images (stimulus duration: 500 ms). We analyzed source-reconstructed data from three ROIs: V1-3, V4t/LO, and IT/PHC. We computed an RDM for each participant, region, and time point. RDMs were computed by extracting an activity pattern for each image and computing pairwise dissimilarities (1 minus Pearson’s r) between the activity patterns. **e)** Schematic overview of model fitting procedure. We tested two model classes: a visuo-semantic model consisting of all category and feature RDMs and a DNN model consisting of all CORnet-Z and CORnet-R layer RDMs. The respective model RDMs serve as predictors. We fitted the two models to the MEG RDMs for each participant, region, and time point, using cross-validated non-negative least squares regression.

For lower-level visual cortex (V1-3), the DNN model explained significant amounts of variance between 60 and 638, and 818 and 884 ms after stimulus onset, while the visuo-semantic model did so between 118 and 660 ms after stimulus onset (Figure 2a). The DNN model explained more variance than the visuo-semantic model during the early (66 - 128 ms) as well as the late (422 - 516 ms, 520 - 544 ms, 820 - 844 ms) phases of the response. For intermediate visual cortex (V4t/LO), the DNN model explained variance between 62 and 610 ms after stimulus onset, while the visuo-semantic model explained variance between 110 and 562 ms after stimulus onset (Figure 2a). The amount of explained variance did not significantly differ between the two models. The results for lower-level visual cortex indicate that the DNN model outperformed the visuo-semantic model at explaining object representations, especially during the early phase of the response (*<* 128 ms after stimulus onset). In contrast, for higher-level visual cortex (IT/PHC), the visuo-semantic model outperformed the DNN model. The DNN model explained variance only between 182 and 270 ms after stimulus onset (Figure 2a). The visuo-semantic model explained variance during a longer time window, between 96 and 658 ms after stimulus onset (Figure 2a). Furthermore, the visuo-semantic model explained more variance than the DNN model between 146 and 488 ms after stimulus onset (specifically 146 - 188 ms, 196 - 234 ms, 326 - 344 ms, 348 - 402 ms, 412 - 464 ms, 468 - 488 ms). In summary, the results across the ventral stream regions show a reversal in which model best explains variance in the RDM movies, from the DNN model in lower-level visual cortex to the visuo-semantic model in higher-level visual cortex.

**Fig. 2.**
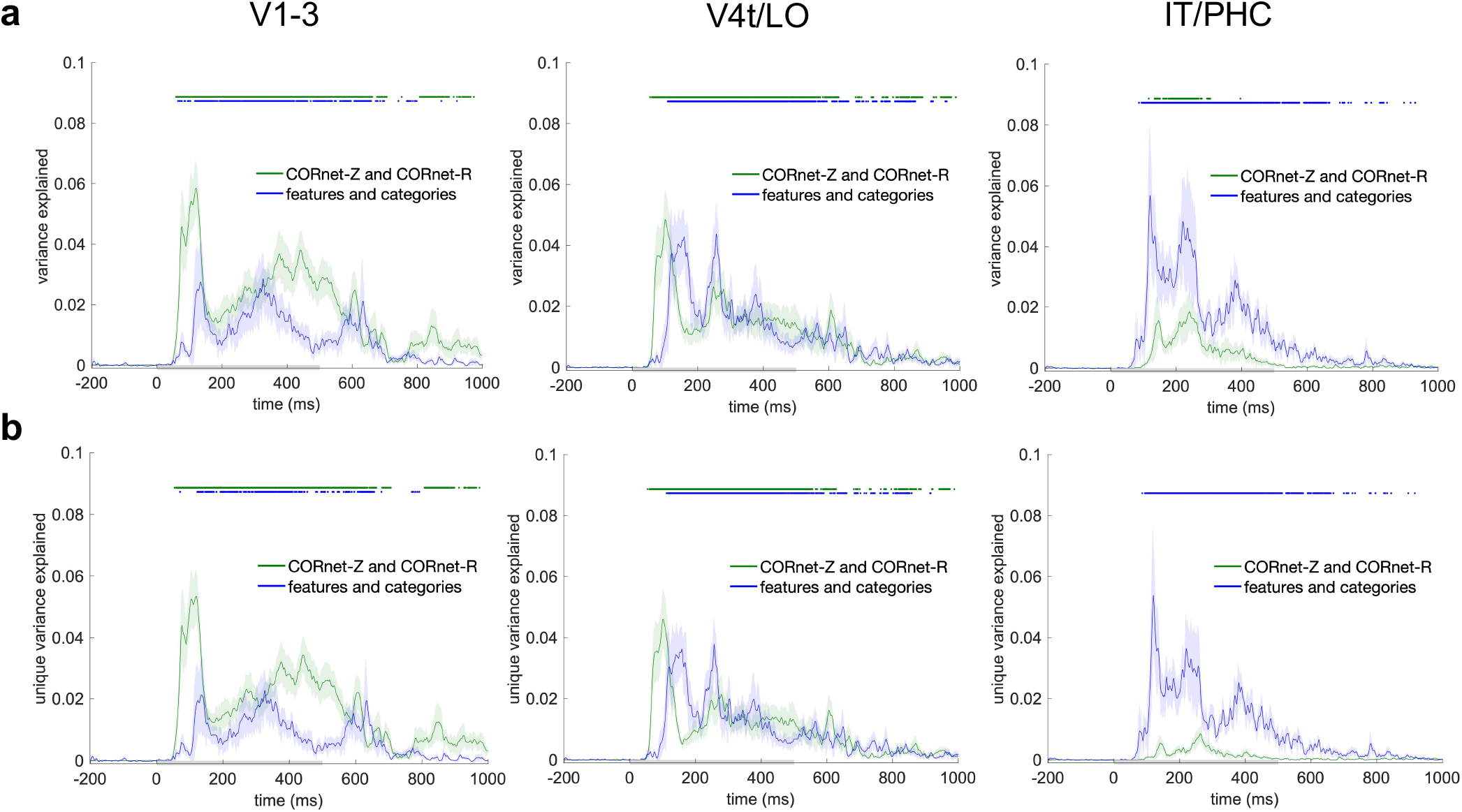
DNNs better explain lower-level visual representations, visuo-semantic models better explain higher-level visual representations. a) Variance explained by the visuo-semantic (blue) and DNN (green) models in the source-reconstructed MEG data. To estimate the variance explained by each model class, we took the model components (visuo-semantic features or DNN layers) and ran a non-negative least squares regression. Variance explained was computed as the variance explained by the model predictions in data for images left out during fitting. Significant variance explained is indicated by a horizontal line above the graph (one-sided Wilcoxon signed-rank test, p < 0.05 corrected). The shaded area around the lines shows the standard error of the mean across participants. The x axis shows time relative to stimulus onset. The gray horizontal bar on the x axis indicates the stimulus duration. **b)** Unique variance explained by the visuo-semantic and DNN models in the source-reconstructed MEG data. To estimate the unique variance explained by each model class, we took the model predictions (one RDM movie per model class, also used in panel a) and ran a second-level non-negative least squares regression. Unique variance explained was computed by subtracting the variance explained by the reduced GLM (excluding the model class of interest) from the total variance explained by the full GLM (including both model classes). Conventions are the same as in panel a.

### Visuo-semantic models explain unique variance in higher-level visual representations

Our results suggest that DNNs and visuo-semantic models explain complementary components of human ventral-stream representational dynamics. To explicitly test this hypothesis, we assessed the unique contributions of the two models. For this, we first computed the best RDM predictions for each model class, and then used the resulting cross-validated RDM predictions in a second-level general linear model (GLM) in which we combined the two model classes. We computed the unique contribution of a model class by subtracting the variance explained by the reduced model (i.e. the GLM without the model class of interest) from the variance explained by the full model (including both model classes). For lower-level visual cortex (V1-3), the DNN model explained unique variance between 60 and 638, and 818 and 884 ms after stimulus onset, while the visuo-semantic model did so between 124 and 412 ms after stimulus onset (Figure 2b). For intermediate visual cortex (V4t/LO), the DNN model explained unique variance between 62 and 610 ms after stimulus onset, while the visuo-semantic model did so between 118 and 546 ms after stimulus onset (Figure 2b). These results indicate that the DNN and visuo-semantic models each explained a significant amount of unique variance in lower-level and intermediate visual cortex compared to the baseline period. However, for lower-level visual cortex, the DNN model explained more unique variance than the visuo-semantic model during the early (66 - 128 ms) as well as the late phases of the response (422 - 516 ms, 520 - 544 ms, 820 - 844 ms). For intermediate visual cortex, the unique variance explained did not significantly differ between the two models. For higher-level visual cortex (IT/PHC), only the visuo-semantic model explained unique variance, between 104 and 640 ms after stimulus onset (specifically 104 - 464 ms, 468 - 500, 542 - 578, and 608 - 640). Furthermore, the visuo-semantic model explained significantly more unique variance than the DNN model between 146 and 488 ms after stimulus onset (specifically 146 - 188 ms, 196 - 234 ms, 326 - 344 ms, 348 - 402 ms, 412 - 464 ms, 468 - 488 ms, Figure 2b). These results indicate that, in the context of a visuo-semantic predictor, the tested DNNs explain unique variance at lower-level but not higher-level stages of visual processing which instead show a unique contribution of visuo-semantic models. Visuo-semantic models appear to explain components of the higher-level visual representations that DNNs fail to fully capture.

### Object parts and basic categories contribute to the unique variance explained by visuo-semantic models in higher-level visual representations

To better understand which components of the visuo-semantic model contribute to explaining unique variance in the higher-level visual representations, we repeated our analyses separately for subsets of object features and subsets of categories. We grouped the visuo-semantic labels into the following subsets: object parts, color, shape, and texture, and subordinate, basic, and superordinate categories (Figure 1b). We found that, among the categories, subordinate and basic categories explained variance in higher-level visual cortex (IT/PHC) (Figure 3a). Furthermore, each of these models explained unique variance in higher-level visual cortex, while the DNN model did not (Figure 3b). Among the object features, only object parts explained variance in higher-level visual cortex (Figure 3a). Furthermore, object parts explained unique variance in higher-level visual cortex, while the DNN model did not (Figure 3b). We next evaluated the three best predictors among the object features and categories together in the context of the DNN predictor. While object parts, subordinate categories, basic categories, and DNNs all explained variance in higher-level visual cortex, only object parts and basic categories explained unique variance (Figure 3b).

**Fig. 3.**
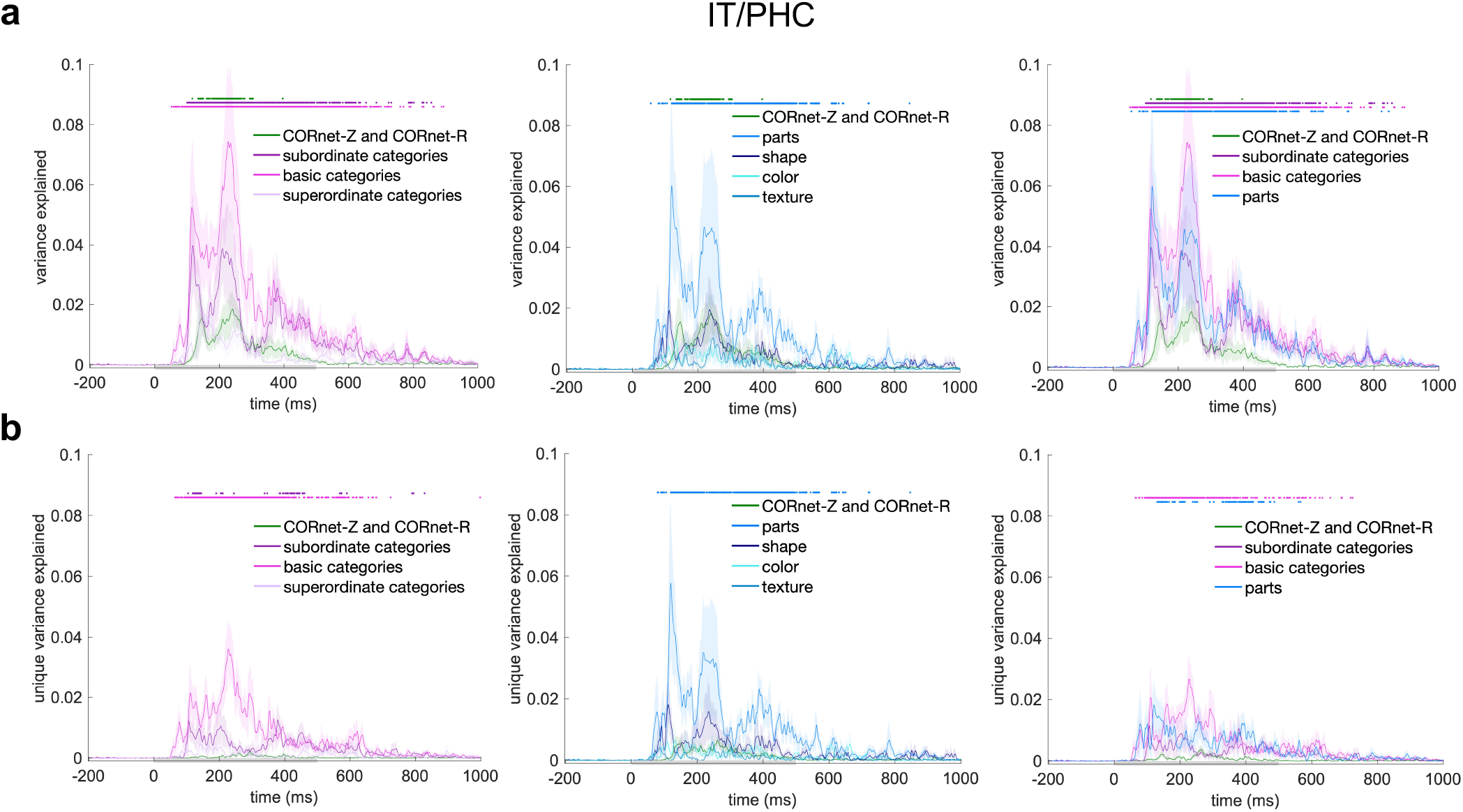
Object parts and basic categories contribute to the unique variance explained by visuo-semantic models in higher-level visual representations. **a)** Variance explained by the object features (parts, shape, color, texture; shades of blue), categories (subordinate, basic, superordinate; shades of pink), and DNN model (green) in the source-reconstructed MEG data. Conventions are the same as in Figure 2a. **b)** Unique variance explained by the features, categories, and DNN model in the source-reconstructed MEG data. Conventions are the same as in Figure 2b.

### Deep neural networks and visuo-semantic models explain complementary components of human ventral-stream representational dynamics

To summarize our results, we computed a model difference score by subtracting the unique variance explained by the visuo-semantic model from that explained by the DNN model in the dynamic ventral-stream representations. Figure 4 displays the model difference score for each of the three regions of interest for the first 300 ms of image processing. Results show a gradual reversal in the relative importance of DNN versus visuo-semantic features in explaining ventral-stream object representations as they unfold over space and time. DNN features lead in lower-level visual areas V1-3 early in time (*<* 128 ms after stimulus onset), while visuo-semantic features lead in higher-level visual areas IT/PHC later in time (*>* 146 ms after stimulus onset). Among the visuo-semantic features, object parts and basic categories drive the advantage over DNNs. While the relative contributions of DNN and visuo-semantic features did not significantly differ in intermediate visual areas V4t/LO, the observed timing of events across the three regions of interest provides complementary information. Within the first 100 ms after stimulus onset, the relative contribution of DNN features peaks in lower-level and intermediate visual areas. Around 120 ms after stimulus onset, the relative contribution of visuo-semantic features peaks in *higher-level* visual areas. Around 150 ms after stimulus onset, the relative contribution of visuo-semantic features peaks in *intermediate* visual areas, following a period of feedback information flow from higher-level to intermediate visual areas (10). Together, these results suggest that DNNs and visuo-semantic models explain complementary components of human ventral-stream representational dynamics. Moreover, our results show that a significant component of the variance unexplained by DNNs in higher-level visual areas is structured, and can be explained by relatively simple, readily nameable aspects of the images.

**Fig. 4.**
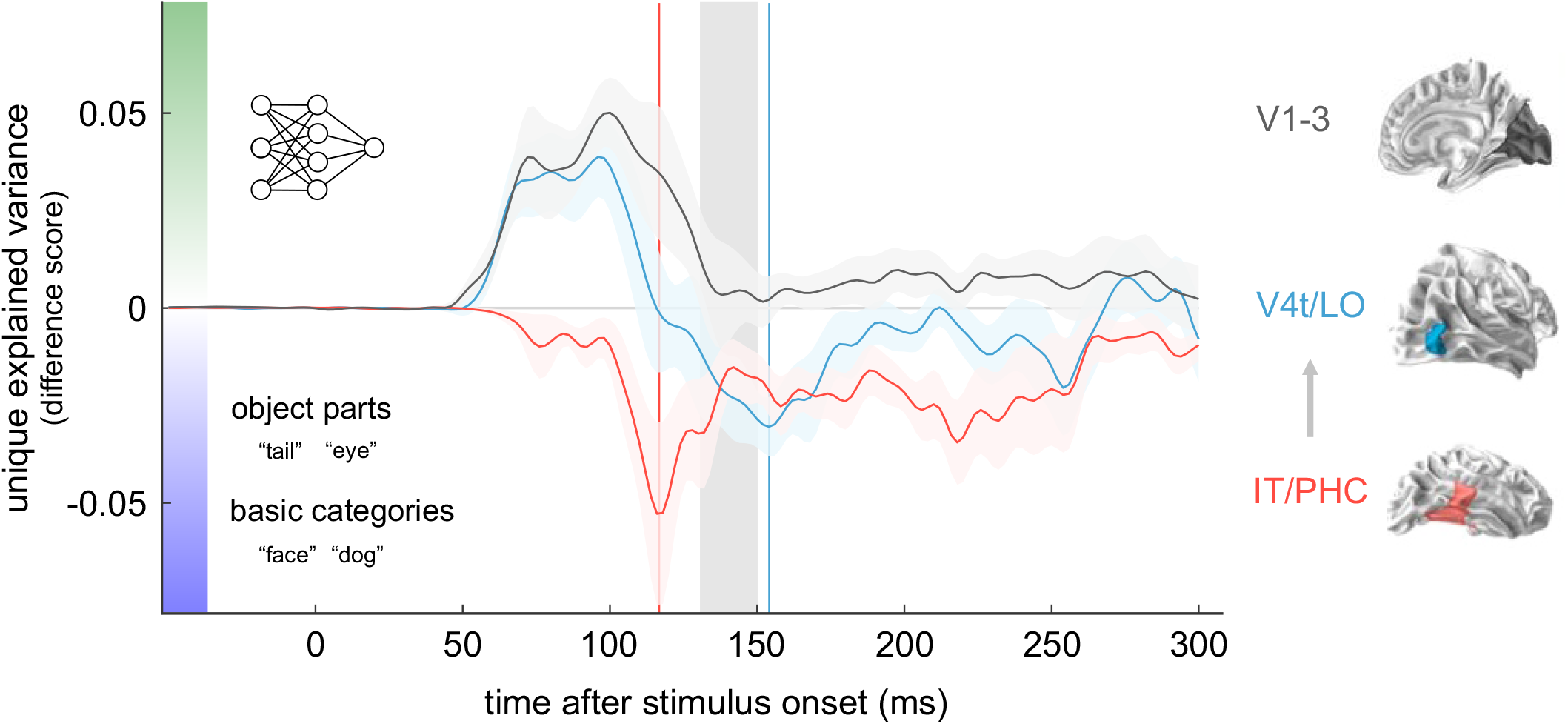
Summary of findings: deep neural networks and visuo-semantic models explain complementary components of human ventral-stream representational dynamics. Between 66 and 128 ms after stimulus onset, DNNs outperform visuo-semantic models in lower-level areas V1-3 (grey line, positive deflection). This early time window is thought to be dominated by feedforward and local recurrent processing. In contrast, starting 146 ms after stimulus onset, visuo-semantic models outperform DNNs in higher-level visual areas IT/PHC (red line, negative deflection). This result shows that DNNs fail to account for a significant component of variance in higher-level cortical dynamics, which is instead accounted for by visuo-semantic features, in particular object parts and basic categories. The peak of visuo-semantic model performance in higher-level areas (red vertical line) precedes the peak in intermediate areas (blue vertical line). This sequence of events aligns with the timing of feedback information flow from higher-level to intermediate areas (light grey rectangle and arrow) as reported in (10). The shaded area around the lines shows the standard error of the mean across participants.

## Discussion

Neural representations of visual objects dynamically unfold over time as we are making sense of the visual world around us. These representational dynamics are thought to reflect the cortical computations that support human object recognition. Here we show that DNNs and human-derived visuo-semantic models explain complementary components of representational dynamics in the human ventral visual stream, estimated via source-reconstructed MEG data. We report a gradual reversal in the importance of DNN and visuo-semantic features from lower-to higher-level visual areas. DNN features explain variance over and above visuo-semantic features in lower-level visual areas V1-3 especially during the early phase of the response (*<* 128 ms after stimulus onset). In contrast, visuo-semantic features explain variance over and above DNN features in higher-level visual areas IT/PHC during a later time window (starting at 146 ms after stimulus onset). Among the visuo-semantic features, object parts and basic categories drive the explanatory advantage over DNNs. Consistent with our hypothesis, our findings suggest that current DNNs fail to fully capture the visuo-semantic features represented in higher-level human visual cortex, and suggest a path towards more accurate models of ventral stream computations.

Our finding that DNNs outperform visuo-semantic models at explaining lower-level cortical dynamics replicates and extends prior functional magnetic resonance imaging (fMRI) work, which showed that DNNs explain response variance across all stages of the ventral stream while visuo-semantic models predominantly explain response variance in higher-level visual cortex (13, 14, 17, 43, 49). However, the contribution of such models to explaining representational dynamics has not been investigated. Using source-reconstructed MEG data, we show that the advantage of DNNs over visuo-semantic models is strongest within the first 128 ms after stimulus onset. During this early time window, the response is likely dominated by feedforward and local recurrent processing as opposed to top-down feedback signals from higher-level areas (7, 11). Our results show the importance of analyzing temporally resolved neuroimaging data for revealing when in time competing models account for the rapid dynamic unfolding of human ventral-stream representations.

Our findings show that DNNs, despite reaching human-level performance on large-scale object recognition tasks (20), fail to fully capture visuo-semantic features represented in higher-level human visual cortex, in particular object parts and basic categories. These results suggest that current DNNs do not fully develop the representational features that give rise to the categorical divisions observed in higher-level visual representations (34–42, 44, 45). In line with this, prior fMRI work showed that DNNs only adequately accounted for higher-level visual representations after adding new representational features (13, 22, 49, 50). The new features were either explicit semantic features (49) or were created by linearly combining DNN features to emphasize categorical divisions observed in the higher-level visual representations, including the division between faces and nonfaces and between animate and inanimate objects (13, 50). Our results are also consistent with an earlier MEG study which showed that adding semantic features to a simpler HMAX model was beneficial for modeling object representations in visual cortex starting around 200 ms after stimulus onset (51). DNNs may, at least in part, use different object features for object recognition than humans do. This conclusion is consistent with prior reports that DNNs rely more strongly on lower-level image features such as texture for object categorization (52).

Our study makes several important contributions to the existing body of work on modeling ventral-stream computations with DNNs. First, our results suggest that introducing locally recurrent connections to DNNs, to more closely match the architecture of the ventral visual stream, is not sufficient to fully capture the representational dynamics observed in higher-level human visual cortex. Second, our results tie together space and time through analysis of source-reconstructed MEG data. We show that DNNs outperform visuo-semantic models in lower-level visual areas V1-3 during the first 128 ms of processing, while visuo-semantic models outperform DNNs in higher-level visual areas IT/PHC starting at 146 ms after image onset. Third, we show that a significant component of the unexplained variance in higher-level cortical dynamics is structured, and can be explained by readily nameable aspects of object images, specifically object parts and basic categories. In prior behavioral work using the same image set and visuo-semantic labels, we showed that category labels, but not object parts, outperformed DNNs at explaining object similarity judgements (48). This highlights a difference with the current results, which show a role for both object parts and categories in accounting for response variance in higher-level visual cortex over and above DNNs. These results are consistent with the idea that, compared to responses in ventral visual cortex, behavioral similarity judgements may more strongly emphasize semantic object information (18, 48, 53). Future studies should extend this work to richer stimulus and model sets.

To build more accurate models of human ventral stream computations, we need to provide DNNs with a more human-like learning experience. Two important areas for improvement are visual diet and learning objectives. Each of these shape the internal object representations that develop during visual learning. Humans have a rich visual diet and learn to distinguish between ecologically relevant categories at multiple levels of abstraction, including faces, humans, and animals (45, 53). DNNs have a more constrained visual diet and are trained on category divisions that do not entirely match the ones that humans learn in the real world. For example, the most common large-scale image dataset for training DNNs with category supervision (13–15, 20, 22, 27, 33, 50, 54), the ILSVRC 2012 dataset (54), contains subordinate categories that most humans would not be able to distinguish, including dog breeds such as “schipperke” and “groenendael”, and lacks some higher-level categories relevant to humans, including “face” and “animal”. The path forward is unfolding along two main directions. The first is enrichment of the visual diet of DNNs by better matching the visual variability present in the real world, for example by increasing variability in viewpoint or by training on videos instead of static images (55, 56). The second is to more closely match human learning objectives, for example by introducing more human-like category objectives or unsupervised objectives (57–60). Training DNNs on more human-like visual diets and learning objectives may give rise to representational features that more closely match the visuo-semantic features represented in human higher-level visual cortex.

## Methods

### Stimuli

Stimuli were 92 colored images of real-world objects spanning a range of categories, including humans, non-human animals, natural objects, and manmade objects (12 human body parts, 12 human faces, 12 animal bodies, 12 animal faces, 23 natural objects, and 21 manmade objects). Objects were segmented from their backgrounds (Figure 1a) and presented to human participants and models on a gray background.

### Visuo-semantic models

Feature and category visuo-semantic models have been described in (45, 48), where further details can be found.

#### Definition of visuo-semantic models

To create visuo-semantic models, human observers generated feature labels (e.g., “eye”) and category labels (e.g., “animal”) for the 92 images (45). The visuo-semantic models are schematically represented in Figure 1b. Feature labels were divided into object parts, colors, textures and shapes, while category labels were divided into subordinate categories, basic categories and superordinate categories. Labels were obtained in a set of two experiments. In Experiment 1, a group of 15 human observers (mean age=26 years; 11 females) generated feature and category labels for the object images. Human observers were native English speakers and had normal or corrected-to-normal vision. In the instruction, we defined features as “visible elements of the shown object, including object parts, colors, textures and shapes”. We defined a category as “a group of objects that the shown object is an example of”. The instruction contained two example images, not part of the 92 object-image set, with category and feature descriptions. We asked human volunteers to list a minimum of five descriptions, both for categories and for features. The 92 images were shown, in random order, on a computer screen using a web-based implementation, with text boxes next to each image for human observers to type category or feature descriptions. We subsequently selected, for categories and features separately, those descriptions that were generated by at least three out of 15 human observers. This threshold corresponds to the number of human observers that, on average, mentioned a particular category or feature for a particular image. The threshold is relatively lenient, but it allows the inclusion of a rich set of descriptions, which were further pruned in Experiment 2. We subsequently removed descriptions that were either inconsistent with the instructions or redundant. Observers generated 212 feature labels and 197 category labels. These labels are the model dimensions. In Experiment 2, a separate group of 14 human observers (mean age=28 years; seven females) judged the applicability of each model dimension to each image, thereby validating the dimensions generated in Experiment 1, and providing, for each image, its value (present or absent) on each of the dimensions. Human observers were native English speakers and had normal or corrected-to-normal vision. During the experiment, the object images and the descriptions, each in random order, were shown on a computer screen using a web-based implementation. The object images formed a column, while the descriptions formed a row; together they defined a matrix with one entry, or checkbox, for each possible image-description pair. We asked the human observers to judge for each description, whether it correctly described each object image, and if so, to tick the associated checkbox. The image values on the validated model dimensions define the model (when agreed by at least 75% of human observers from Experiment 2). To increase the stability of the models during subsequent fitting, we iteratively merged binary vectors that were highly correlated (r > 0.9), alternately computing pairwise correlations between the vectors, and averaging highly-correlated vector pairs, until all pairwise correlations were below threshold. The final full feature and category models consisted of 119 and 110 dimensions, respectively.

#### Construction of the visuo-semantic representational dissimilarity matrices

To compare the models to the measured brain representations, the models and the data should reside in the same representational space. This motivates transforming our models to RDM space. For each model dimension, we computed, for each pair of images, the squared difference between their values on that dimension. The squared difference reflects the dissimilarity between the two images in a pair. Given that a specific feature or category can either be present or absent in a particular image, image dissimilarities along a single model dimension are binary: they are zero if a feature or category is present or absent in both images, and one if a feature or category is present in one image but absent in the other. The dissimilarities were stored in an RDM, yielding as many RDMs as model dimensions. The full feature RDM model consists of 119 RDMs; the full category RDM model consists of 110 RDMs. Subsets of the feature and category RDM models consist of the following number of RDMs: object parts (82), shape (15), color (10), texture (12), subordinate categories (38), basic categories (67), superordinate categories (5).

### Deep neural networks

CORnet-Z and CORnet-R architectures have been described in (27), where further details can be found.

#### Architecture and training

We used feedforward (CORnet-Z) and locally recurrent (CORnet-R) (27) models in our analyses. The architectures of the two DNNs are schematically represented in Figure 1b. The architecture of CORnets is inspired by the anatomy of monkey visual cortex where each processing stage in the model is thought to correspond to areas V1, V2, V4, and IT respectively (27). The output of the last model area is mapped to the model’s behavioral choices using a linear decoder. We chose these DNNs because they have similar architectures with one being feedforward and the other locally recurrent, their architecture is inspired by the primate visual system, they are one of the best models for predicting visual responses in human IT (32, 33) and monkey IT (20), and their architectures are relatively simple compared to other DNNs. Each “visual area” in CORnet-Z (“Zero”) consists of a single convolution, followed by a ReLU nonlinearity and max pooling. CORnet-R (“Recurrent”) introduces local recurrent dynamics within an area. The recurrence occurs only within an area; there are no bypass or feedback connections between areas. For each area, the input is down-scaled twofold and the number of channels is increased twofold by passing the input through a convolution, followed by group normalization (61) and a ReLU nonlinearity. The area’s internal state (initially zero) is added to the result and passed through another convolution, again followed by group normalization and a ReLU nonlinearity, resulting in the new internal state of the area. At time step 0 “t0” there is no input to “V2” and above layers, and as a consequence no image-elicited activity is present. From time step “t1” onwards, the image-elicited activity is present in all “visual areas” as the output of the previous area is immediately propagated forward. CORnet-R was trained using five time steps (“t0” - “t4”). Both DNNs were trained on 1.2 million images from the ILSVRC data base (54). The task was to classify each image as containing an object in one of 1,000 possible categories.

#### Construction of the DNN representational dissimilarity matrices

Representations of the 92 images were computed from the layers of CORnet-Z and CORnet-R. For CORnet-Z, we included the decoder layer and the final processing stage (output) from each “visual area” layer, which resulted in 5 layers. For CORnet-R, we included the decoder layer and the final processing stage from each “visual area” layer for each time step, which resulted in 21 layers. For each layer of CORnet-Z and CORnet-R, we extracted the unit activations in response to the images and converted these into one activation vector per image. For each pair of images, we computed the dissimilarity (1 minus Spearman’s correlation) between the activation vectors. This yielded an RDM for each DNN layer. The resulting RDMs capture which stimulus information is emphasized and which is de-emphasized by the DNNs at different stages of processing.

### MEG source-reconstructed data

Acquisition and analysis of the MEG data have been described in (6), where further details can be found. The source reconstruction of the MEG data has been described in (10), where further details can be found.

#### Participants

Sixteen healthy human volunteers participated in the MEG experiment (mean age = 26, 10 females). MEG source reconstruction analyses were performed for a subset of 15 participants for whom structural and functional MRI data were acquired. Participants had normal or corrected-to-normal vision. Before scanning, the participants received information about the procedure of the experiment and gave their written informed consent for participating. The experiment was conducted in accordance with the Ethics Committee of the Massachusetts Institute of Technology Institutional Review Board and the Declaration of Helsinki.

#### Experimental design and task

Stimuli were presented at the centre of the screen for 500 ms, while participants performed a paper clip detection task. Stimuli were overlaid with a light gray fixation cross and displayed at a width of 2.9° visual angle. Participants completed 10 to 14 runs. Each image was presented twice in every run in random order. Participants were asked to press a button and blink their eyes in response to a paper clip image shown randomly every 3 to 5 trials. These trials were excluded from further analyses. Each participant completed two MEG sessions.

#### MEG data acquisition and preprocessing

MEG signals were acquired from 306 channels (204 planar gradiometers, 102 magnetometers) using an Elekta Neuromag TRIUX system (Elekta) at a sampling rate of 1,000 Hz. The data were bandpass filtered between 0.03 and 330 Hz, cleaned using spatiotemporal filtering, and down-sampled to 500 Hz. Baseline correction was performed using a time window of 100 ms before stimulus onset.

#### MEG source reconstruction

The source reconstructions were performed using the MNE Python toolbox (62). We used participant individual structural T1 scans to obtain volume conduction estimates using single layer boundary element models (BEMs) based on the inner skull boundary. Instead of BEMs being based on the FreeSurfer watershed algorithm originally used in the MNE Python toolbox, we extracted BEMs using FieldTrip as the original method yielded poor reconstruction results. The source space consisted of 10,242 source points per hemisphere. The source points were positioned along the gray/white matter boundary, as estimated via FreeSurfer. We defined source orientations as surface normals with a loose orientation constraint. We used an iterative closest point procedure for MEG/MRI alignment based on fiducials and digitizer points along the head surface, after initial alignment based on fiducials. We estimated the sensor noise covariance matrix from the baseline period (100 ms to 0 ms before stimulus onset) and regularized it according to the Ledoit–Wolf procedure (63). We projected source activations onto the surface normal, obtaining one activation estimate per point in source space and time. Source reconstruction allowed us to estimate temporal dynamics in specific brain regions. Source reconstruction provides an estimate of what brain regions the information is coming from rather than a direct measurement of representations in different brain regions (see (64) for a discussion on that topic).

#### Definition of regions of interest

We used a multimodal brain atlas (65) to define ROIs. We defined three ROIs covering low-level (V1–3), intermediate (V4t/LO1–3), and high-level visual areas (IT/PHC, consisting of TE1-2p, FFC, VVC, VMV2–3, PHA1–3). We converted the atlas annotation files to fsaverage coordinates (66) and mapped them to each participant using spherical averaging.

#### Construction of the MEG representational dissimilarity matrices

We computed temporally changing RDM movies from the source-reconstructed MEG data for each participant, ROI, hemisphere, and session. We first extracted a trial-average multivariate source time series for each stimulus. We then computed an RDM at each time point by estimating the pattern distance between all pairs of images using correlation distance (1 minus Pearson correlation). The RDM movies were averaged across hemispheres and sessions, resulting in one RDM movie for each participant and ROI.

### Weighted representational modeling

Weighted representational modeling used here has been described in (13, 45, 48, 50, 67), where further details can be found. We could predict the brain representations by making the assumption that each model dimension contributes equally to the representation. We use the squared Euclidean distance as our representational dissimilarity measure, which is the sum across dimensions of the squared response difference for a given pair of stimuli. The squared differences simply sum across dimensions, so the model prediction would be the sum of the single-dimension model RDMs. However, we expect that not all model dimensions contribute equally to brain representations. To improve model performance, we linearly combined the different model dimensions (features and categories for the visuo-semantic models, layers for the DNNs, see below for details) to yield an object representation that best predicts the source-reconstructed MEG data. Because the squared differences sum across dimensions in the squared Euclidean distance, weighting the dimensions and computing the RDM is equivalent to a weighted sum of the single-dimension RDMs. When a dimension is multiplied by weight *w*, then the squared differences along that dimension are multiplied by *w*^*2*^. We can therefore perform the fitting on the RDMs. We estimated the representational weights, one for each single-dimension RDM, using regularized (L2) linear regression, implemented in MATLAB using Glmnet (https://hastie.su.domains/glmnet_matlab/?). Glmnet implements elastic net regularized regression using cyclical coordinate descent. We used standard settings (including standardization of the predictors before fitting), except that we constrained the weights to be non-negative. To prevent biased model performance estimates due to overfitting to a particular set of images, model performance was estimated by cross-validation to a subset of the images held out during model fitting. For each cross-validation fold, we randomly selected 84 of the 92 images as the training set and eight images as the test set (with the constraint that test images had to contain four animate objects (two faces and two body parts) and four inanimate objects) and used the corresponding pairwise dissimilarities to estimate the model weights. The model weights were then used to predict the pairwise dissimilarities for the eight left-out images. This procedure was repeated many times until predictions were obtained for all pairwise dissimilarities. For each cross-validation fold (using different test and train images each time), we determined the best regularization parameter (i.e. the one with the minimum squared error between prediction and data) using nested cross-validation to held-out images within the training set. We performed the fitting procedure for each participant, each time point, and each brain region of the MEG source-reconstructed data.

### Variance and unique variance analysis

We used a GLM to evaluate variance and unique variance explained by the models in the source-reconstructed MEG data (10). For each model m, the variance explained *R*^2^ was computed by GLM fitting using X = “model m” and Y = data. For each model m, the unique variance explained was computed by subtracting the total variance explained by the reduced GLM (excluding the model of interest) from the total variance explained by the full GLM. Specifically, for model m, we fit the GLM using X = “all models but m” and Y = data, then we subtracted the resulting *R*^2^ from the total *R*^2^ (fit the GLM using X = “all models” and Y = data). We performed this procedure for each participant and ROI. We used non-negative least squares to find optimal weights as RDMs consist of dissimilarity estimates, which cannot be negative. A constant term was included in the GLM model to correct for homogeneous changes in dissimilarity across the whole RDM. To evaluate the significance of the variance explained and unique variance explained by each model across participants, we first subtracted an estimate of the prestimulus baseline in each participant and then performed a one-sided Wilcoxon signed-rank test against 0. The prestimulus baseline was defined as the average (unique) variance explained between 200 - 0 ms before stimulus onset. We also tested if and when the (unique) variance explained differed between the two models using a two-sided Wilcoxon signed-rank test. We controlled the expected false discovery rate at 0.05 across time points for each model evaluation, model comparison, and ROI. Variance and unique variance explained and standard error lines were low-pass filtered at 80 Hz (Butterworth IIR filter; order 6) for easier visibility. Statistical inference is based on unsmoothed data.

## ACKNOWLEDGEMENTS

This research was supported by the Wellcome Trust grant [206521/Z/17/Z] awarded to KMJ; the Alexander von Humboldt Foundation postdoctoral fellowship awarded to KMJ, the German Research Council grants [CI241/1-1, CI241/3-1 CI241/7-1] awarded to RMC, the European Research Council grant [ERC-StG-2018-803370] awarded to RMC, and a Natural Sciences and Engineering Research Council of Canada Discovery Grant [RGPIN-2019-06741] awarded to MM. We thank Martin Schrimpf for giving his input on the CORnets Methods section. For the purpose of open access, the author has applied a CC BY public copyright licence to any Author Accepted Manuscript version arising from this submission.

## AUTHOR CONTRIBUTIONS

KMJ, TCK, RMC, NK, and MM designed the experiments. KMJ collected behavioral data for visuo-semantic model building. RMC collected MEG data. TCK provided source-reconstructed MEG data and second-level GLM code. KMJ performed the analyses. KMJ and MM wrote the paper. All authors edited the paper. NK and MM supervised the work.

## DATA AVAILABILITY

The datasets generated during the current study are available from the corresponding authors on request.

## CODE AVAILABILITY

The code generated during the current study is available from the corresponding authors on request.

